# Cannabidiol (CBD) as a novel inhibitor of HLA-G expression in human choriocarcinoma cell line (JEG-3)

**DOI:** 10.1101/2025.07.02.662795

**Authors:** Kevin I. Martínez, María B. Palma, Fernando J. Sepúlveda, Damián E. Moavro, Edgardo D. Carosella, Marcela N. García, Fernando L. Riccillo

**Affiliations:** Cátedra de Citología, Histología y Embriología, Facultad de Ciencias Médicas, Universidad Nacional de La Plata, La Plata, Argentina; Consejo Nacional de Investigaciones Científicas y Técnicas (CONICET); LIAN‑CONICET, Fundación FLENI, Buenos Aires, Argentina; Laboratorio de Biología Celular, Departamento de Biología Celular, Universidad de Concepción, Concepción, Chile; Atomic Energy and Alternative Energies Agency (CEA), Hematology and Immunology Research Division, Saint-Louis Hospital, Paris, France; U976 HIPI Unit, University of Paris, Paris, France; Cátedra Histología y Embriología Animal, Facultad de Ciencias Naturales y Museo, Universidad Nacional de La Plata, La Plata, Argentina

**Keywords:** HLA-G, CBD, high-CBD extract, JEG-3, immunomodulation

## Abstract

Cannabinoids have emerged as promising agents in cancer research due to their antitumor properties. While their effects on tumor growth and survival have been widely investigated, their impact on immune checkpoint regulation remains largely unexplored. In this study, we examined the effects of cannabidiol (CBD) and a high-CBD extract (CBD-HCE) on the expression of HLA-G in human choriocarcinoma JEG-3 cells. HLA-G is a non-classical HLA class I molecule associated with immune escape in tumors. Both CBD and CBD-HCE were found to reduce cell proliferation and migration, increase apoptosis, and significantly downregulate HLA-G expression at both the mRNA and protein levels. This inhibitory effect was observed to be both dose- and time-dependent, and it was completely reversible after the treatment was removed, indicating a dynamic and CBD-dependent modulation. These results provide the first experimental evidence of HLA-G downregulation by CBD and CBD-HCE, highlighting a novel immunomodulatory mechanism with potential therapeutic implications. By simultaneously targeting tumor cell viability and immune evasion, CBD-based compounds may enhance antitumor immune responses and improve the efficacy of immunotherapies. Further research involving additional tumor cell lines, in vivo models, and immune-relevant systems are necessary to validate and expand upon these findings.

## INTRODUCTION

The discovery of cannabinoid (CB) receptors in the early 1990s sparked renewed interest in cannabis research, leading to the elucidation of the endocannabinoid system (ECS). The ECS is highly conserved throughout evolution and dates back at least 550 million years [1, 2]. It comprises a diverse family of molecules, their receptors, and the associated proteins responsible for their metabolism, including synthesis, transport, and degradation. Subsequently, an extensive body of research has emerged, shedding light on the (patho) physiological roles of the ECS [3, 4, 5, 6, 7].

Cannabinoids are chemical compounds belonging to the group of terpenophenols, which exert their action from their association with specific membrane receptors [8, 9, 10, 11]. They are classified into three groups: a) phytocannabinoids (natural cannabinoids of plant origin, from the plant C. sativa); b) synthetic cannabinoids and c) endogenous cannabinoids (endocannabinoids): N-arachidonoylethanolamine (anandamide) (AEA) and 2-arachidonoylglycerol (2-AG). The two most abundant phytocannabinoids and those best characterized by their therapeutic effects are Δ9-tetrahydrocannabinol (THC) and Cannabidiol (CBD). THC is the main psychoactive agent of cannabis but also has analgesic, anti-inflammatory, antiemetic and orexigenic properties [12, 13, 14]. On the other hand, CBD modulates the psychotropic effects of THC, has antipsychotic, neuroprotective, immunomodulatory, and anti-inflammatory, antitumor, antidiabetic and other properties, such as the ability to reduce tobacco addiction [12, 13, 14, 15]. However, the action of several of the other cannabinoids, as well as their acid forms, should not be underestimated.

Phytocannabinoids exert their effects by mimicking the action of endogenous cannabinoids, especially through their specific CB1 and CB2 receptors [10, 11]. Currently, other alternative receptors have been described (especially for CBD, given its low affinity for CB1 and CB2) such as orphan G protein-coupled receptors (GPR55, GPR19, GPR18), the transient receptor potential cation channel subfamily V type 1 and 4 (TRPV1, TRPV4), the serotonin 5-HT1A receptor, the γ aminobutyric acid (GABA) receptor and the γ peroxisome proliferator-activated receptor (PPARγ) [16,17].

Cannabinoids have clearly been shown to exert a palliative effect in cancer patients. (4,12, 18, 19). However, the therapeutic potential of cannabinoids in oncology is not restricted to their use as palliative care agents. A great number of studies have shown that THC, CBD and other cannabinoids exhibit antitumor effects in a wide range of *in vitro* and *in vivo* cancer models [4, 19, 20, 21, 22, 23]. Nevertheless, the specific mechanisms through which cannabinoids modulate tumor biology are not yet fully understood. This is partly due to their pleiotropic mechanisms of action, as they can act through multiple receptors. Furthermore, it is extremely important to consider that the effects of cannabinoids depend on the concentrations, types, and/or combinations of cannabinoids involved, as well as the specific tumor cells they target. The HLA-G is a non-classical class I HLA molecule which, along with other isotypes, plays a key role in feto-maternal tolerance. It offers protection to the fetus by shielding it from the maternal immune system, thereby preventing its rejection [24]. In addition, it was observed that HLA-G contributes to allogeneic tissue graft tolerance [25, 26, 27]. Whether membrane-bound or soluble, HLA-G exhibits strong binding affinity to its inhibitory receptors present on immune cells such as NK cells, T and B cells, and monocytes/dendritic cells. This binding leads to the inhibition of effector functions, resulting in immune suppression and promoting a tolerogenic environment.

On the other hand, HLA-G-expressing tumors can exploit its immunosuppressive properties to evade immune surveillance, primarily through direct interaction with various immune effectors [28, 29, 30]. This interaction plays a significant role in modulating immune responses and promoting tumor survival.

Recent studies have shown that THC may impair the therapeutic efficacy of PD-1 blockade by acting on CB2 receptors expressed on tumor-specific T cells [31]. In contrast, other reports have provided novel evidence supporting the immunostimulatory effects of CBD, including its ability to upregulate MHC-I expression [32] and enhance the inhibition of the PD-1 immune checkpoint [33]. Nevertheless, no data are currently available regarding the potential modulatory role of CBD—or other cannabinoids—on the expression or function of the immune checkpoint HLA-G. To investigate this potential relationship, we selected the human choriocarcinoma cell line (JEG-3) as our in vitro model, given its high and constitutive expression of HLA-G, providing a consistent baseline that improves sensitivity for the detection of potential cannabinoid-induced changes. This type of cancer that occurs exclusively in the female population is one of the most aggressive gestational trophoblastic tumors with a high propensity for metastasis [34, 35]

This study aimed to explore, for the first time, the potential relationship between cannabinoids and HLA-G expression in a tumor cell line (JEG-3). Furthermore, it provides the first characterization of cannabinoid-induced effects on key parameters of cell proliferation and death in this human choriocarcinoma cell model.

## MATERIALS AND METHODS

### Chemicals

Cannabidiol (CBD, purity by HPLC: 99.8%) and *Cannabis sativa* extract with high content in cannabidiol (i.e., high-CBD content extract: CBD-HCE) were used.

Cannabis sativa extract was supplied by Plan Cannabis Civil Association Foundation (File No. 119264/22-1, Legal Entities, Ministry of Justice, Buenos Aires, Argentina). Extraction was obtained using ethanol and then evaporated. The cannabinoid profiles of the extract were quantified against commercial THC, CBD, CBN and CBG standards (Cerilliant^©^ - Texas,USA) by HPLC/UV-DAD (Shimadzu LC-20A) School of Medical Science, National University of La Plata. For CBD-HCE composition see Table 1.

**Table 1.** Content of the main phytocannabinoids in high-cannabidiol (CBD) content extract (CBD-HCE)

| Phytocannabinoid | Content (mg/ml) |
| --- | --- |
| Cannabidiol acid (CBDA) | 0.523 |
| Cannabidiol (CBD) | 15.579 |
| Cannabigerol (CBG) | 0.355 |
| Cannabinol (CBN) | 0.032 |
| Tetrahydrocannabinol acid (THCA) | 0.208 |
| Tetrahydrocannabinol (THC) | 0.770 |

The CBD was purchased from Cerilliant^©^ (Texas, USA). It was initially dissolved in dimethyl sulfoxide (DMSO) to a concentration of 250 mM and stored at -20°C. CBD was further diluted with tissue culture medium for in vitro studies, keeping the DMSO concentration below 0.2 %.

### Cell culture

JEG-3 human choriocarcinoma cell line was used. It was generously provided by Instituto de Fisicoquímica Biológica y Química, Universidad de Bioquímica y Farmacia (UBA-CONICET), Buenos Aires.

This cell line was cultured in vitro in Dulbecco’s modified Eagle’s medium (DMEM) supplemented with 10% foetal bovine serum (Gibco^TM^) according with the manufacturer’s protocols and 1% penicillin/streptomycin (Gibco^TM^) in a 5% CO2, humidified atmosphere at 37 °C until a confluence state of 75%. Cells were regularly dissociated using Trypsine-EDTA 0.25% for further immunocytochemistry (ICC) and reverse transcription quantitative polymerase chain reaction (RT-qPCR) studies.

### Cell viability assay (MTT)

To assess the impact of CBD and CBD-HCE on cell viability and their antiproliferative effect, we conducted the MTT colorimetric assay using [3-(4,5-dimethyl-2-thiazolyl)-2,5-diphenyl-2H tetrazolium bromide] obtained from Santa Cruz Biotech, USA. JEG-3 cells were seeded (1 x10^4^ cells/well) in a 96-well flat-bottom multiwell plate (Corning Inc.® USA), in 100 µL of DMEM with 0.4 % DMSO.

After 24 h., cells were treated with CBD and CBD-HCE at different concentrations (0, 1, 2.5, 5, 10, 20, 40, 80, 100, 150 µM) during 24 h. Following incubation with the two cannabinoid treatments, MTT (0.5 mg/ml final concentration) was added to each well and maintained for 3 h. The insoluble formazan crystals were solubilized by the addition of 200 ml/well of 100% DMSO, and the optical density (OD) was measured using an automatic microplate reader (Beckman Coulter DTX 880 Microplate Reader, Fullerton, CA, USA) at 560 nm with 640 nm as the reference wavelength. The IC_50_ values of each extract were determined from the concentration-effect curves (% cell death) using non-linear regression analysis.

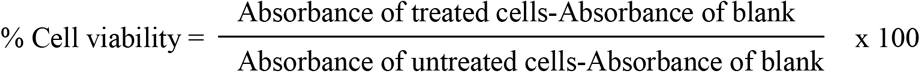

### Immunocytochemistry

For immunocytochemistry, 2 × 10^6^ cells were seeded on a silanized glass in cell culture medium for 24 h. The medium was then removed and replaced with the corresponding treatment solutions: 1 µM or 5 µM of CBD or CBD-HCE, for 24/48 h. After treatment, samples were washed with phosphate-buffered saline (PBS, pH 7.4), and the cells were fixed with PATHOFIX® (Biopack). Following fixation, a second PBS wash was performed, and endogenous peroxidase activity was blocked with 3% H_2_O_2_ for 10 minutes. Cells were then incubated overnight with 1:50 dilutions of the following primary monoclonal antibodies (Santa Cruz Biotech, CA, USA): anti-HLA-G, anti-Ki-67, anti-cleaved caspase-3, and anti-MMP-9. Detection was performed using the ABC system (Vector, USA) with diaminobenzidine as the chromogen. Finally, the samples were lightly counterstained with Mayer’s hematoxylin. Positive and negative controls were included in each step to validate staining specificity.

### Cell morphometry

Morphometric analyses were performed using ImageJ software (v1.54d, NIH, Bethesda, MD, USA) on images previously captured and digitized with the Micrometrics LE system (NY, USA). Cell proliferation was assessed by calculating: (1) the percentage of cells in mitosis (M phase) relative to the total number of cells per field (% MF = [mitotic cells/total cells] x 100); and (2) the percentage of Ki-67–positive nuclei (% Ki+ = [Ki-67^+^ cells/total cells] x 100). For each treatment condition, three independent experiments were conducted. In each case, 15 non-overlapping high-power fields (400X magnification) were analyzed, and the average of these 15 measurements was obtained. The mean ± SEM of the values obtained for each of the three experimental replicates was then calculated. Apoptotic activity was evaluated by quantifying the Caspase-3 immunostained area (IA Cas-3), calculated as the ratio between the DAB-stained area and the corresponding reference area. Migratory capacity was assessed similarly, using the MMP-9–positive stained area (IA MMP-9). In both cases, the reference area corresponded to each high-power field at 400X magnification. As above, 15 fields were analyzed per experiment, and the averaged values were used to calculate the mean ± SEM for each treatment.

### Cell migration assay

In a 6-well plate, 1 × 10^6^ cells/well were seeded in growth medium until reaching approximately 80% confluence. The culture medium was then removed, and a solution of DMEM + 10% FBS + 0.4% DMSO was used for the control group, while 1 µM CBD or 1 µM CBD-HCE was added for the treatment groups. Both control and treatment solutions were renewed every 12 hours. Eight independent replicates were analyzed for each condition (N = 8). For each replicate, a 150 µm-wide gap was created at time zero (*t*_*0*_), and wound closure was monitored at 24, 30, and 40 hours post-injury. The percentage of wound closure (regenerated area, % AR) was calculated using the formula: % AR = [(Gap area at t_0_ – Gap area at t_t_) / Gap area at t_0_] × 100, where *t*_*0*_ is the initial time and *t*_*t*_ corresponds to each evaluation time point. For each condition and time point, the values obtained from the eight replicates were averaged, and results are expressed as mean ± SEM. Cell proliferation, migration and death rates were also evaluated throughout this assay.

### CBD and CBD-HCE Treatments for HLA-G Expression Analysis

#### Incubation assay

In a 6 multi-well plate, 5 × 10^5^ cells/well were seeded in cell culture medium for 24 h. After removing the culture medium, a cell sample was removed as an initial control (t = 0). Then, the remaining wells were incubated with a solution of DMEM + 10% FBS + 0.4% DMSO as a control medium, or with the following treatments: 1 µM or 5 µM of CBD or CBD-HCE for 12 h, 24 h and 36 h. Finally, the cells were isolated and treated for analysis of gene expression levels by RT-qPCR.

#### Reversal assay

In a 6 multi-well plate, 5 × 10^5^ cells/well were seeded in cell growth medium for 24 h. The culture medium was then removed, and the cells were incubated for 36 h with 1 µM CBD. Finally, the solution was removed, cells were washed with PBS, and fresh growth medium was added for 24-and 36-h post-treatment incubation. At each point, a cell sample was removed for subsequent mRNA extraction and HLA-G expression analysis by RT-qPCR to test for possible recovery of HLA-G basal expression.

### RNA Extraction, cDNA Synthesis, and RT-qPCR Analysis

For the analysis of HLA-G expression in tumor cells cultured under the different experimental conditions, reverse transcription-polymerase chain reaction (RT-PCR) was performed using specific primers to detect all known isoforms, published previously (Tronik-Le Roux D et al., 2017). RNA extraction from JEG-3 cells, was performed with TRIzol Reagent (Invitrogen). For cDNA synthesis, 500–1000 ng of the total RNA was retro-transcribed with MMLV reverse transcriptase (Promega), according to manufacturer’s instructions. For HLA-G detection by RT-qPCR, cDNA samples were diluted fivefold, and it was performed with StepOne Plus Real Time PCR System (Applied Biosystems). The FastStart Universal SYBR Green Master Mix (Roche) was used for all reactions. Primers efficiency and initial molecule (N_0_) values were determined by LinReg software 3.0, and gene expression was normalized to RPL7 housekeeping gene, for each condition. All the oligonucleotide sequences are listed in Table 2.

**Table 2.**
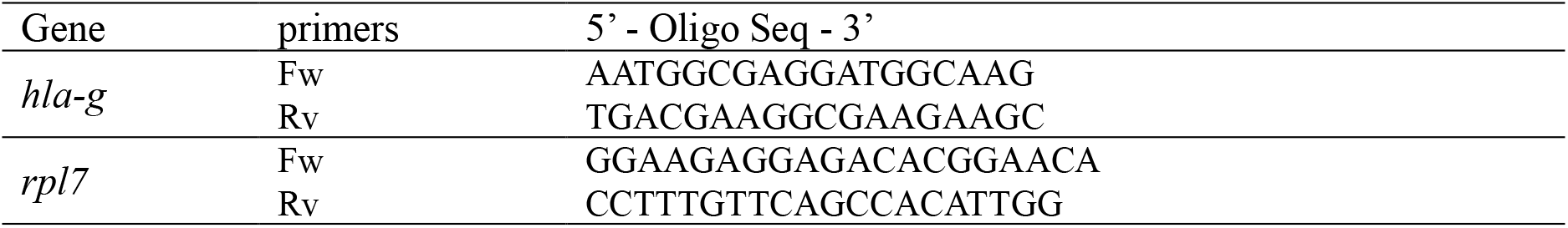
Oligonucleotide sequences of *hla-g* gene and *rpl7* reference gene.

### Statistics

Significant differences were determined using Graph Pad Prism 8 (USA). Statistical significance was calculated using t-tests and ANOVA. Tukey-Kramer post hoc analyses were conducted when appropriate. The significance was set at p < 0.05. Data were expressed as mean ± standard error of mean (SEM).

## RESULTS

### Determination of Non-Cytotoxic Concentrations by MTT Assay

The MTT assay established the tolerable cannabinoid concentrations for cells under different treatment and incubation conditions. Fig. 1 shows the viability curve for each treatment, which exhibited similar IC_50_ values (CBD = 18.78 µM; CBD-HCE = 18.27 µM). The values shown represent the average of all measurements performed (n=10) for each treatment. Based on these data, we observed that cell viability was not significantly affected below 7,5 µM. Therefore, we selected 1 µM and 5 µM as safe working concentrations for subsequent experiments.

**Figure 1.**
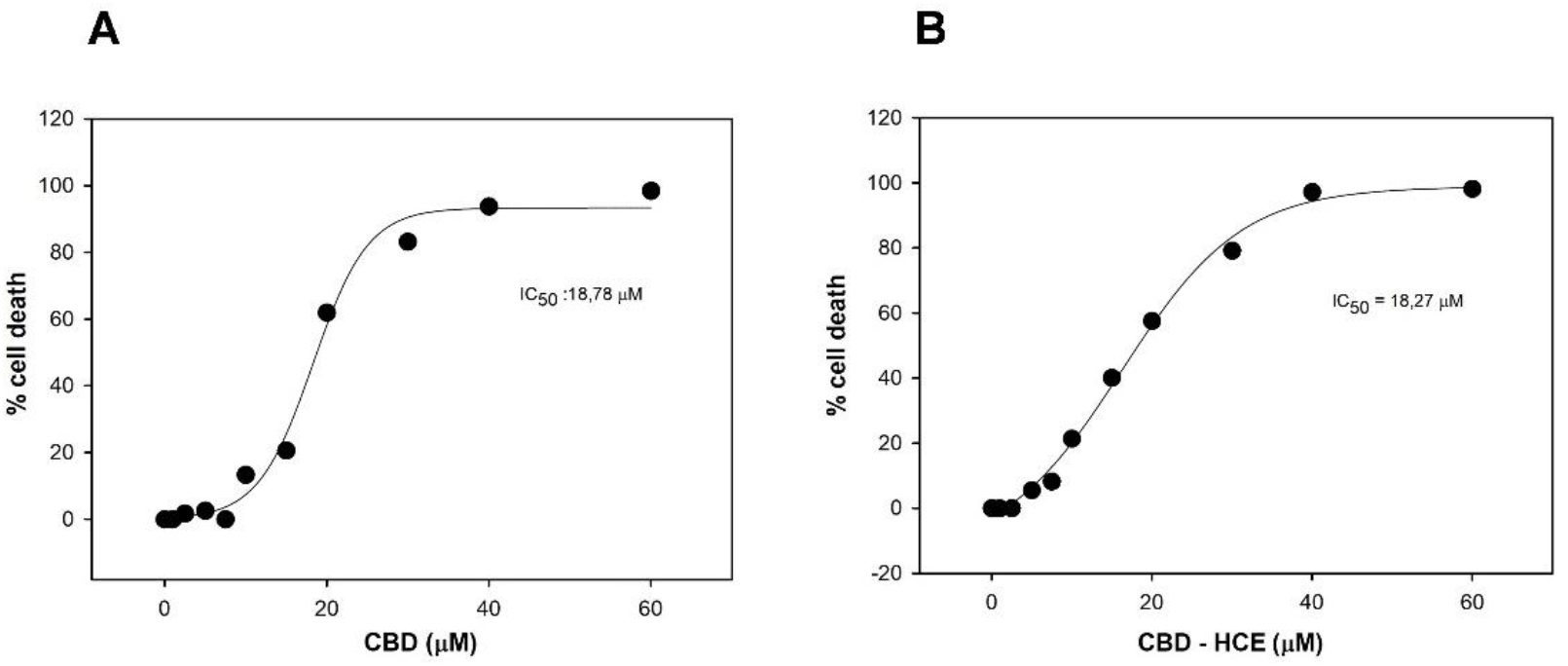
Cell viability assay (MTT). (a) Effect of cannabidiol (CBD, 0–60 µM) and (b) high-CBD extract (CBD-HCE, 0–60 µM) on the viability of JEG-3 cells after 24-hour exposure. Cell viability was assessed using the MTT assay to determine non-cytotoxic working concentrations. Results are expressed as percentage of cell death relative to cannabinoid concentration. The half-maximal inhibitory concentration (IC_50_) was calculated for each treatment (CBD: 18.8 µM; CBD-HCE: 18.27 µM). Each point represents an average of ten independent measurements (n = 10).

### CBD and CBD-HCE induce cell death of human choriocarcinoma cancer cells

The analysis of the pro-apoptotic effect of CBD and CBD-HCE in JEG-3 cells by Cas-3 labeling, showed significantly higher values (p<0,001) in both treatments: CBD (1 µM: 0.23 ± 0.003 y 5 µM: 0.24 ± 0.007) and CBD-HCE (1 µM: 0.23 ± 0.002 y 5 µM: 0.24 ± 0.006) compared to the values observed in control cultures (0.15± 0,02). No significant differences were observed between the two cannabinoid treatments (p > 0,05). (Fig. 2a and 2c).

**Figure 2.**
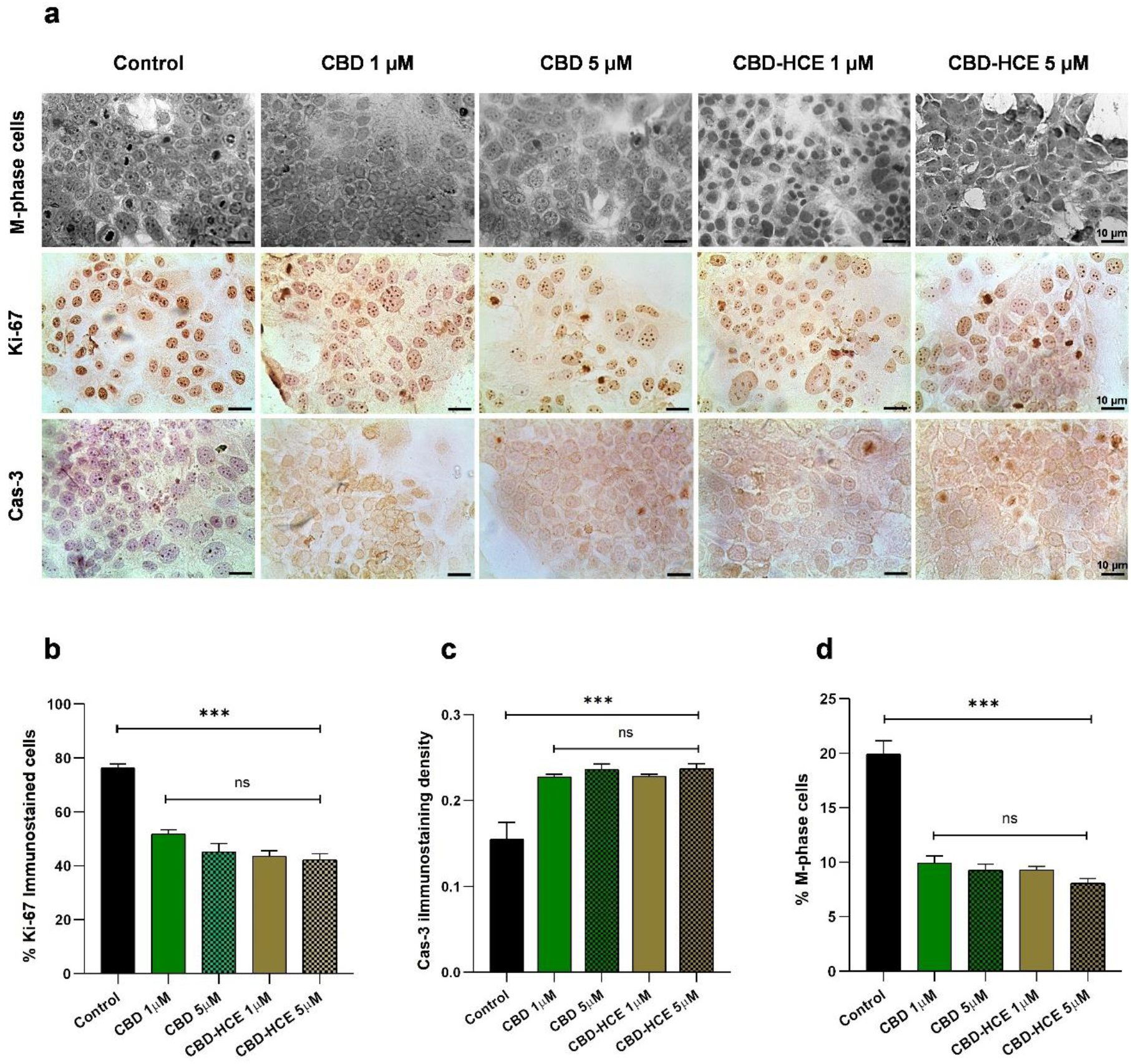
Characterization of the antitumor effects of CBD and CBD-HCE on JEG-3 cells. (a) Representative micrographs showing the effects of 1 µM and 5 µM CBD and CBD-HCE on JEG-3 cells. The top row displays cells in M phase, the middle row shows Ki-67–positive cells, and the bottom row corresponds to Caspase-3–immunolabeled cells. Scale bar = 5 µm. (b) Quantification of Ki-67–positive nuclei (%), demonstrating a reduction in proliferation upon treatment. (c) Caspase-3 labeling density, showing a significant increase in apoptotic activity in treated cells. (d) Percentage of cells in M phase under each treatment condition. Solid bars: 1 µM; dotted bars: 5 µM. Each bar represents the mean ± SEM of three independent experiments (n = 3). ***p < 0.001.

### CBD and CBD-HCE inhibit cell proliferation of human choriocarcinoma cancer cells

Proliferation analysis using Ki-67 immunolabeling showed a significant reduction (p < 0.001) in the percentage of proliferating cells following treatment with CBD (1 µM: 51.78% ± 1.96; 5 µM: 45.47% ± 3.71) and CBD-HCE (1 µM: 43.32% ± 2.45; 5 µM: 42.17% ± 2.76) compared to the control (76.23% ± 1.76). No significant differences were observed between CBD and CBD-HCE at the same concentrations (p > 0,05). (Fig. 2a and 2b).

Consistent with the Ki-67 data, mitotic index analysis also revealed a significantly higher percentage of cells in M phase in the control group (19.93% ± 1.05) compared to those treated with CBD (1 µM: 9.92% ± 0.57; 5 µM: 9.27% ± 0.47) and CBD-HCE (1 µM: 9.29% ± 0.27; 5 µM: 8.07% ± 0.38) (p < 0.001). (Fig. 2a and 2d).

### CBD and CBD-HCE inhibit cell migration of human choriocarcinoma cancer cells

Following monolayer injury, the percentage of wound closure after 24 hours was significantly higher in the control group (60.30 % ± 5.50; p < 0.01) compared to cultures treated with 1 µM CBD (39.90 % ± 3.90) or 1 µM CBD-HCE (41.60 % ± 3.60) (Fig. 3a and 3b). Each value represents the mean ± SEM of eight independent replicates (n = 8) per treatment. Subsequent measurements at 30 h and 40 h revealed a progressive reduction in wound area in all conditions (Fig. 3b, table 3). While control cultures achieved complete confluence by 40 h, treated cultures exhibited a slower regenerative response, suggesting that achieving complete closure would require additional culture time (Fig. 3b)

**Table 3.** Migration assay. Percentages (%) of area recovery after injury monolayer in control and treated cultures during 24, 36 and 40 h. Data represent the mean ± SEM (n = 8).

|  |  | Time (h) |  |  |
| --- | --- | --- | --- | --- |
|  |  | 24 | 30 | 40 |
| Regenerated Area (%) | Control | $60.30 \pm 5.50$ | $87.82 \pm 5.654$ | $100 \pm 0.000$ |
| | CBD $1\mu\text{M}$ | $9.90 \pm 3.90$ | $66.42 \pm 3.422$ | $92.84 \pm 2.545$ |
| | CBD-HCE $1\mu\text{M}$ | $1.60 \pm 3.60$ | $67.46 \pm 2.186$ | $98.91 \pm 2.133$ |

**Figure 3.**
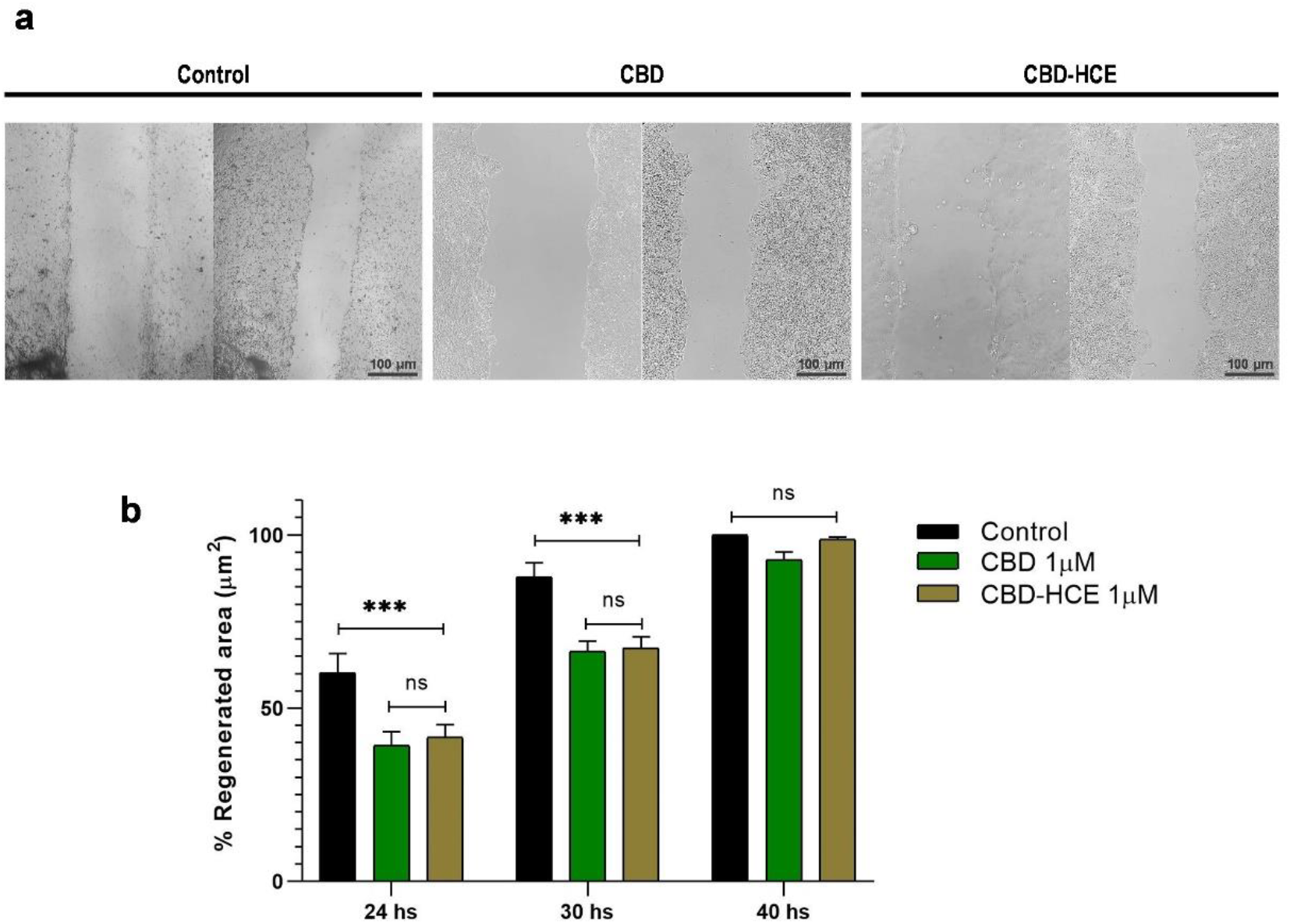
Analysis of the antimigratory effect of CBD and CBD-HCE on JEG-3 cells. (a) Representative images of the wound-healing assay at 0 h and 24 h post-injury under control conditions and after treatment with 1 µM CBD or 1 µM CBD-HCE. Scale bar: 50 µm. (b) Quantification of the regenerated area at 24, 30, and 40 h post-injury for each condition. Black bars indicate control values, green bars correspond to CBD treatment, and brown bars to CBD-HCE treatment. Data represent the mean ± SEM of eight independent experiments (n = 8). ***p < 0.001 vs. control.

Assessment of cell proliferation and apoptosis during migration assays revealed a significant reduction in the mitotic index (M phase) at 24 h in cultures treated with CBD (11.44% ± 0.21) and CBD-HCE (9.81% ± 0.65), relative to control cells (23.96% ± 1.50) (Fig. 4a and 4b). Similarly, Ki-67^−^immunolabeled cells were significantly lower (p < 0.001) in the treated cultures (CBD: 42.18% ± 4.21 and CBD-HCE: 37.58% ± 5.24) compared to the control ones (64.20% ± 3.72) (Fig. 4a and 4c). Regarding apoptosis, no significant differences were observed between treated and control cells (p > 0.05; control: 0.030 ± 0.004, CBD: 0.024 ± 0.005, CBD-HCE: 0.024 ± 0.006; n = 15) (Fig. 4a and 4d). An additional parameter associated with cell migration was MMP-9 expression. MMP-9 immunostaining revealed a significant decrease (p < 0.001) in both cannabinoid-treated groups compared to control (0.0079 ± 0.0007), with CBD-HCE-treated cells showing the lowest expression levels (CBD: 0.0043 ± 0.0006; CBD-HCE: 0.0018 ± 0.0007) (Fig. 4a and 4e).

**Figure 4.**
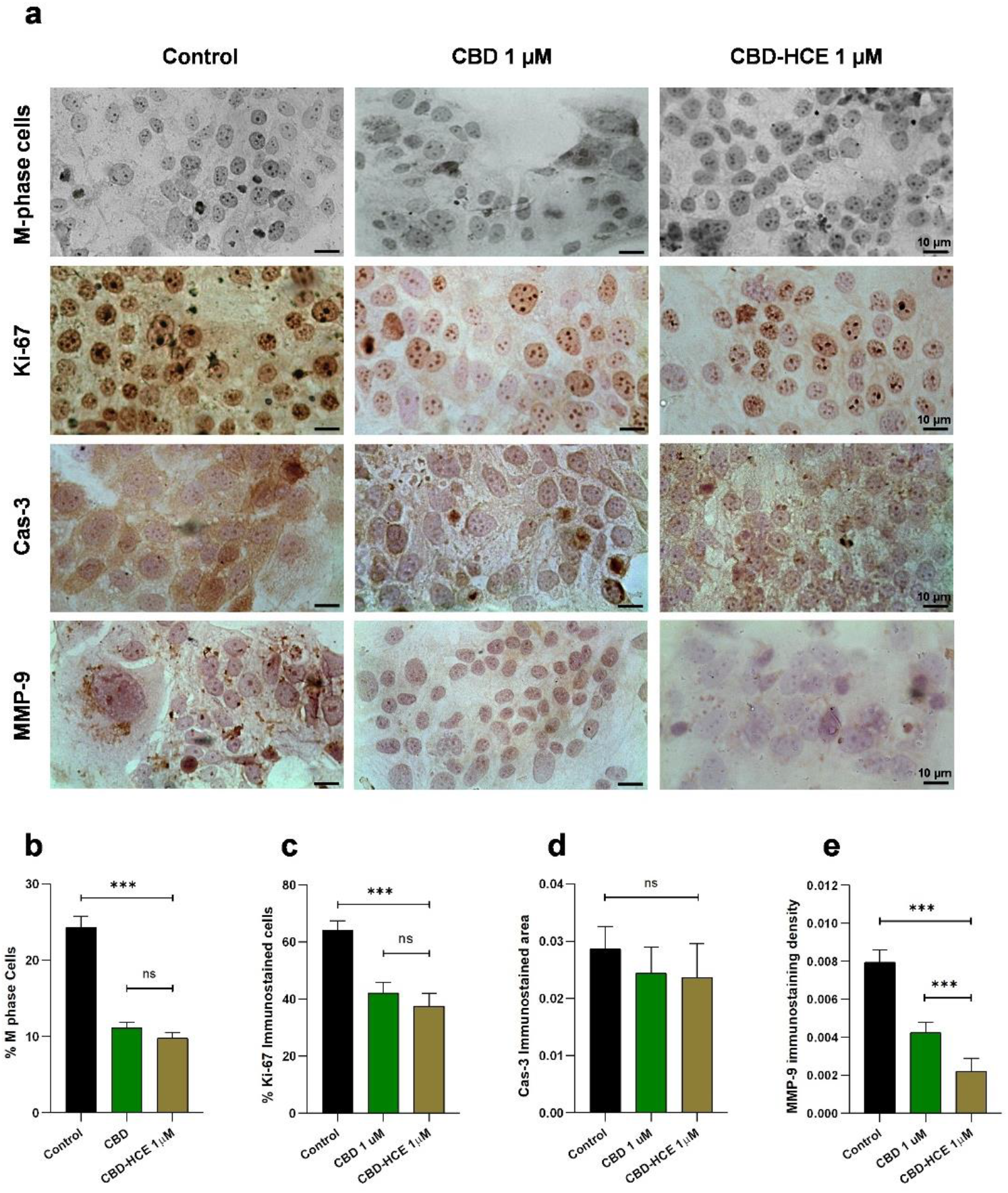
Analysis of proliferation, apoptosis, and migratory capacity in post-injury JEG-3 cells under control conditions or treated with 1 µM CBD or CBD-HCE. (a) Microphotographs of cells in M phase (top row), cells immunostained with anti-Ki-67 mAb (second row), with anti-Caspase-3 mAb (third row), and with anti-MMP-9 mAb (bottom row). Bar = 10 µm. (b) Percentage of cells in M phase. (c) Percentage of Ki-67–positive nuclei. (d) Caspase-3 labeling density. (e) MMP-9 labeling density. Cells treated with CBD and CBD-HCE exhibited reduced proliferation and decreased MMP-9 expression, with no significant changes in apoptosis markers. Black bars represent control values, green bars indicate CBD treatment, and brown bars correspond to CBD-HCE treatment. Each bar shows the mean ± SEM of five independent experiments (n = 5; 15 fields analyzed per treatment). ***p < 0.001.

### CBD and CBD-HCE down-regulate mRNA and protein HLA-G expression

#### RT-qPCR

RT-qPCR analysis revealed a significant decrease in HLA-G mRNA expression following incubation with both 1 µM and 5 µM CBD (Fig. 5a). Expression levels are shown as relative values normalized to the RPL7 reference gene. In the untreated control (t = 0), the HLA-G mRNA level was 0.153 ± 0.006. For both concentrations, mRNA levels were assessed at 12 h, 24 h, and 36 h (Fig. 5b). At 1 µM CBD, HLA-G expression was significantly reduced at all time points compared to the control (p < 0.001), with lower values at 24 h (0.030 ± 0.003) and 36 h (0.028 ± 0.003) compared to 12 h (0.059 ± 0.006). A similar pattern was observed at 5 µM CBD, where expression levels were also significantly lower than control at all time points (p < 0.001): 0.030 ± 0.003 (12 h), 0.023 ± 0.002 (24 h), and 0.037 ± 0.001 (36 h) (n = 8, Fig. 5b).

**Figure 5.**
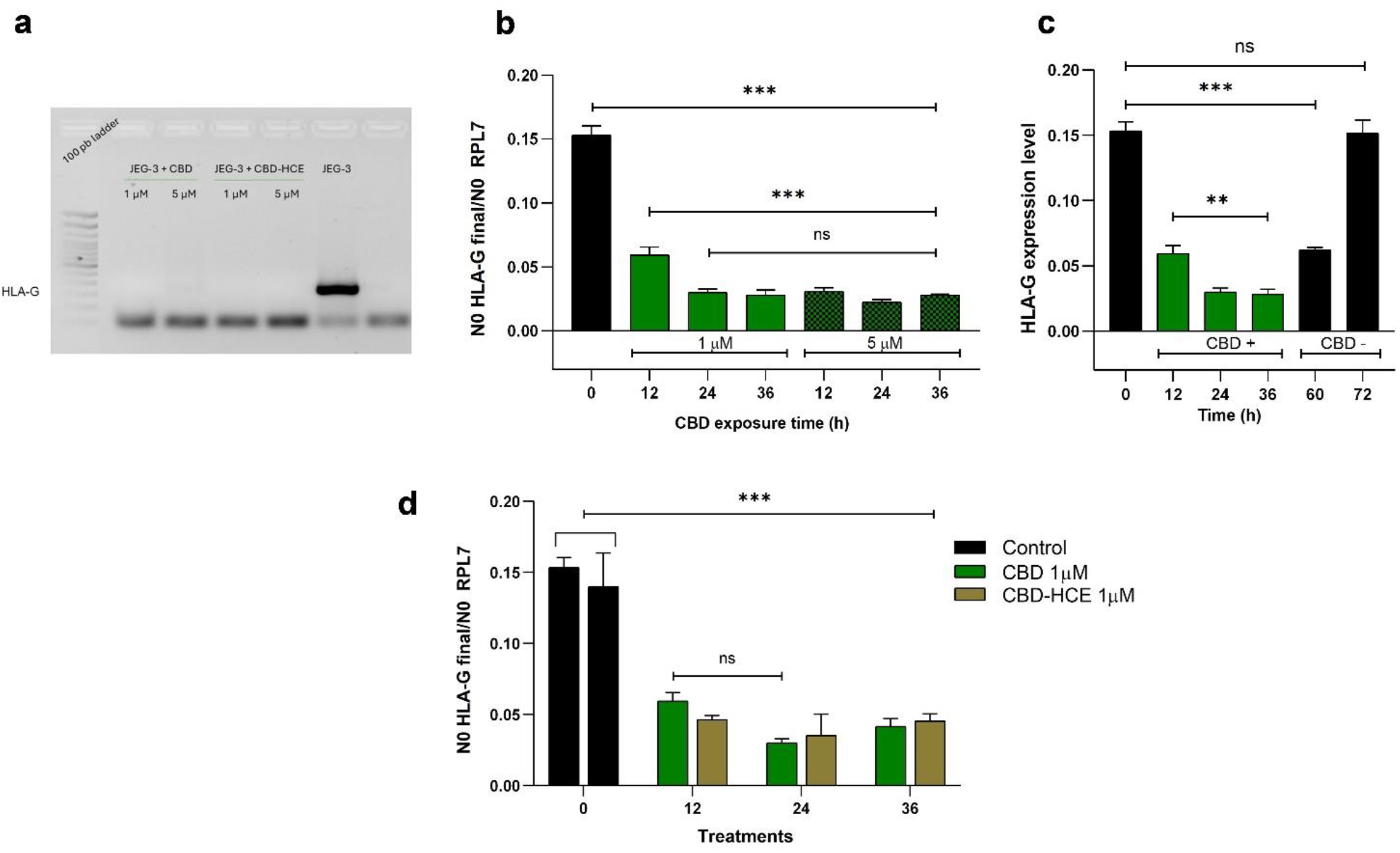
Analysis of HLA-G expression in JEG-3 cells following treatment with CBD and CBD-HCE. (a) Representative agarose gel image showing HLA-G mRNA amplification in untreated control cells and in cells treated for 24 h with either 1 µM or 5 µM of CBD or CBD-HCE. A marked reduction in HLA-G expression is observed in all treated groups. (b) Quantitative RT-PCR analysis of HLA-G expression normalized to RPL7, showing the effect of 1 µM (solid green bars) and 5 µM (dotted green bars) CBD at different time points (12 h, 24 h, 36 h). Untreated control (t = 0 h) is shown in black. (c) Reversibility assay: after 36 h treatment with 1 µM CBD, cells were washed and incubated with fresh medium for an additional 24 h (t_t_ = 60 h) and 36 h (t_t_ = 72 h). HLA-G mRNA expression progressively returned to baseline levels. (d) Comparative qRT-PCR analysis of HLA-G expression after 24 h exposure to 1 µM CBD vs. 1 µM CBD-HCE, indicating similar inhibitory effects. Each bar represents the mean ± SEM of eight independent experiments (n = 8). ***p < 0.001 vs. control.

To further confirm and extend our findings, we investigated whether the inhibitory effect of CBD on HLA-G could be replicated using a high-CBD extract (CBD-HCE) at 1 µM.

Consistently, both treatments at 1 µM led to a significant decrease (p < 0.001) in HLA-G expression compared to the control (CBD-HCE: 12 h: 0.0463 ± 0.003; 24 h: 0.0353 ± 0.01 y 36 h: 0.04548 ± 0.005; control: 0.154 ± 0.01) (Fig. 5d)

#### Immunocytochemistry

Immunocytochemistry analysis revealed clear differences in HLA-G protein levels between treated and control cell cultures (Fig 6a). The quantified immunolabeled area for HLA-G showed a significant reduction in cells treated with CBD at both 1 µM (0.54 ± 0.03) and 5 µM (0.54 ± 0.02) compared to the control, which was normalized to 1 (100%) as the reference value (Fig. 6b). Similarly, treatment with CBD-HCE resulted in a significant decrease (p < 0.001) in HLA-G expression, with values of 0.72 ± 0.07 at 1 µM and 0.38 ± 0.09 at 5 µM (n = 15). Moreover, the reduction observed at 5 µM CBD-HCE was significantly greater than that at 1 µM (p < 0.001) (Fig. 6b).

**Figure 6.**
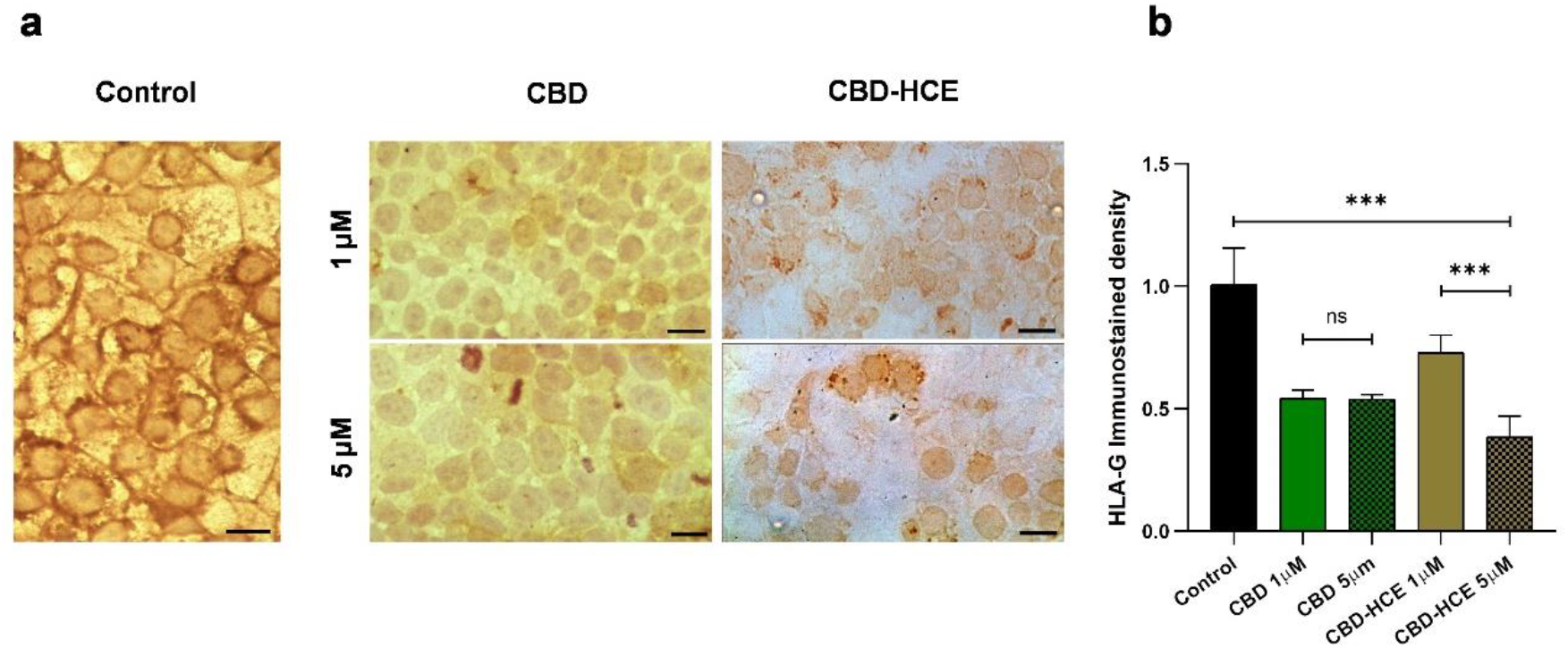
Immunocytochemical analysis of HLA-G expression in JEG-3 cells following CBD and CBD-HCE treatment. (a) Representative light microscopy images of JEG-3 cells immunostained with anti-HLA-G monoclonal antibody under control conditions (Ct) and after treatment with 1 µM or 5 µM of CBD or CBD-HCE. Scale bar = 10 µm. (b) Morphometric quantification of HLA-G expression based on the relative immunolabeled area, expressed as a percentage relative to the control condition (set at 100 %). Bars represent the mean ± SEM of eight independent experiments (n = 8): black (control), green (CBD), and brown (CBD-HCE), with solid and dotted bars indicating 1 µM and 5 µM, respectively. ***p < 0.001.

### HLA-G down-regulation by CBD is reversible

Before treatment (t = 0), HLA-G mRNA expression was 0.153 ± 0.006. As described above and shown in figure 5c, exposure of the cells to 1uM CBD resulted in a significant decrease in mRNA levels. Following CBD removal and replacement with a CBD-free medium, marked expression recovery under drug-free conditions was observed. At 24 h post-removal (t = 60 h), expression levels began to recover (0.062 ± 0.002; p < 0.01), and by 36 h (t = 72 h), had returned to baseline values (0.151 ± 0.009; p < 0.001). Data represent the mean ± SEM from eight independent experiments (n = 8).

## DISCUSSION

Currently, a substantial body of evidence demonstrates the antitumor activity of cannabinoids across multiple cancer types, including lung, breast, ovarian, prostate, pancreas and colon cancers [36, 37, 38, 39, 40, 41]. However, most of the mechanisms underlying these complex and pleiotropic effects remain under extensive research. Most studies have focused on the two main bioactive components of cannabis, THC and CBD, analyzed either as isolated compounds or in standardized extracts. Both phytocannabinoids have shown efficacy in treating several conditions, including cancer. Given the psychotropic side effects of THC, there has been growing interest in CBD, a non-psychoactive cannabinoid. In recent years, its effects and mechanisms of action have been increasingly studied, not only in oncology but also in a wide range of pathological conditions. [23, 42].

Several studies have shown that both CBD and high-CBD extract can inhibit cell proliferation and trigger apoptosis in different tumor cell types [37, 38, 41, 43]. In this study, we focused on analyzing the potential interaction between cannabinoids and HLA-G expression, a non-classical HLA class I molecule involved in immune escape. For the first time, we provide evidence that both CBD and a CBD-HCE extract significantly reduce HLA-G expression in a human choriocarcinoma cell line (JEG-3), which constitutively expresses this immunoregulatory protein. Moreover, we describe for the first time the antitumor properties of these cannabic compounds in JEG-3 cells, a highly aggressive, metastatic trophoblastic tumor model.

To characterize the effect of cannabinoids on this cell line, we analyzed their impact on proliferation, apoptosis, and cell migration. Consistent with previous reports in other *in vitro* and *in vivo* tumor models, we observed a significant reduction in cell proliferation and a marked increase in apoptosis as an outcome of cannabinoid treatments. When assessing the cells’ migratory capacity during monolayer wound healing, pretreatment with CBD or the CBD-HCE impaired their regenerative response. Interestingly, the apoptotic rate during this regenerative phase did not differ significantly from controls, suggesting a possible compensatory mechanism in injured cells, even under the apoptotic-inducing effect of cannabinoids.

Analysis of MMP-9 expression—an essential metalloproteinase involved in extracellular matrix remodeling and cell migration—revealed that both CBD and the CBD-HCE reduced its expression, with the extract exerting a more pronounced inhibitory effect. These results further support the inhibitory role of cannabinoids on tumor cell motility by downregulating molecules critical for invasive behavior.

These findings confirm that JEG-3 cells exhibit similar patterns of proliferation, apoptotic activity and changes in migratory behavior in response to cannabinoid treatments, that is consistent with that observed in other tumor cell lines. Beyond these well-established antitumor effects of cannabinoids, we will focus on discussing their relationship with the expression of HLA-G, a key immune checkpoint molecule involved in the modulation and suppression of immune responses.

Metastatic tumors can subvert the immune system through several mechanisms. One of the most frequently observed in carcinomas is the loss or mutation of genes involved in the MHC-I antigen presentation machinery (APM), rendering tumor cells invisible to cytotoxic T lymphocytes, the key effectors of the adaptive immune response [44, 45]. Another common immune evasion strategy is the upregulation of immune checkpoint (IC) molecules [27]. Recent experimental studies have shown that cannabinoids may counteract metastatic immune escape in vitro, either by enhancing MHC-I surface expression [32] or by reducing the expression of immune-checkpoint molecules such as PD-L1 [33]. In addition, aberrant expressions of non-classical HLAs within the tumor microenvironment (TME) can result from various factors, including complex immunoregulatory signals and intratumoral heterogeneity [46]. This condition further influences the interaction between tumor cells and the immune system, potentially altering the mechanisms of recognition and immune response [47, 48].

This study provides the first evidence of a reversible, cannabinoid-mediated suppression of *HLA-G* expression. Our data demonstrate that both CBD and a CBD-HCE exert a marked inhibitory effect on *HLA-G* mRNA expression in JEG-3 choriocarcinoma cells. This effect was evident after 12 hours of treatment, at both 1 µM and 5 µM concentrations, and persisted thereafter with similar expression patterns for both compounds. Notably, CBD-HCE displayed a comparable inhibitory profile to that of pure CBD. Protein-level analysis by ICQ confirmed the inhibitory effect of both treatments, revealing a decrease in HLA-G immunostaining, with the strongest reduction observed in cells treated with 5 µM of the extract.

As described in the Results section, HLA-G expression was restored to baseline within 36 hours following CBD removal from the culture medium. This reversibility indicates a direct and transient effect, dependent on sustained CBD exposure. The recovery kinetics suggest that CBD targets upstream elements controlling HLA-G transcription or mRNA turnover. To further investigate this mechanism, additional experiments are needed to determine whether this regulation occurs at the transcriptional, post-transcriptional or both levels.

Recent studies have described poor response to immunotherapy in cancer patients using cannabis [49]. In addition, Xiong et al. (2022) demonstrated that both cannabis-derived THC and the endocannabinoid AEA decreased the efficacy of PD-1 blockade by suppressing T-cell-mediated antitumor immune responses by inhibiting JAK/ STAT signaling through CB2 receptor activation. However, these findings are restricted to THC and cannabis-derived THC as the study did not investigate the effects of other cannabinoids. Therefore, its conclusions cannot be extended beyond the specific context of THC-related immunosuppression.

Our results demonstrate a marked antitumor effect after treatment with both pure CBD and a CBD-HCE in JEG-3 cells, inhibiting cell proliferation and cell migration and increasing the rate of apoptosis. Furthermore, the treatment with both pure CBD and a high-CBD extract in JEG-3 cells triggered a marked downregulation of HLA-G expression. These findings are consistent with those reported by Dada et al., 2023 and Sun et al., 2023, reinforcing the potential of CBD as an immunomodulatory compound capable of enhancing, rather than suppressing, host antitumor immune responses, especially in tumors characterized by aberrant immune checkpoint expression.

In summary, our study provides original in vitro evidence that CBD and CBD-rich extracts not only inhibit key tumor cell functions but also consistently downregulate HLA-G expression, a critical immune checkpoint. While these findings are promising, they are limited to a single tumor cell line; thus, further research using additional models and more complex systems is essential to fully elucidate CBD’s immunomodulatory role and therapeutic potential.

## Ethical approval

All experimental procedures were conducted in accordance with relevant guidelines and regulations (the ethical standards of the 1964 Declaration of Helsinki) and were approved by the ethics committee COBIMED (Comite de Bioetica y Etica de la Investigacion de la Facultad de Ciencias Medicas de la Universidad de La Plata).

## Data Availability

The raw data for this study is available from the corresponding author upon reasonable request.

## Acknowledgments

The authors thank Javiera Marini for technical assistance. Also, authors are so grateful to Dra. Analía Seoane, for her selfless collaboration in sharing equipment.

## Author contributions

Project development and experimental designs: F.L.R., K.I.M; Formal analysis: F.L.R., K.I.M., M.B.P., M.N.G., F.J.S.; Methodology: K.I.M, M.B.P., D.E.M.; Resources: F.L.R., M.B.P.,

M.N.G., F.J.S., E.D.C.; Writing-original draft: F.L.R., K.I.M; Writing-review & editing: F.L.R., K.I.M., M.B.P., M.N.G., F.J.S., E.D.C. All authors contributed to the final draft.

